# A conundrum of r-protein stability: unbalanced stoichiometry of r-proteins during stationary phase in *Escherichia coli*

**DOI:** 10.1101/2022.02.09.479841

**Authors:** Kaspar Reier, Petri-Jaan Lahtvee, Aivar Liiv, Jaanus Remme

**Affiliations:** Institute of Molecular and Cell Biology, University of Tartu, Riia 23b 51010, Tartu, Estonia; Department of Chemistry and Biotechnology, Tallinn University of Technology, Akadeemia tee 15 12618, Tallinn, Estonia

## Abstract

Bacterial ribosomes are composed of three ribosomal RNA (rRNA) and over 50 ribosomal protein (r-protein) molecules. R-proteins are essential for ribosome assembly, structural stability, and also participate in almost all ribosome functions. Ribosomal components are present in stoichiometric amounts in the mature 70S ribosomes during the exponential and early stationary growth phases. Ribosomes are degraded in stationary phase, however, the stability and fate of r-proteins during stationary growth phase are not known. In this study, we report a quantitative analysis of ribosomal components during the extended stationary phase in *E.coli*. We show that: (a) the quantity of ribosomes per cell mass decreases in the stationary phase. (b) 70S ribosomes contain r-proteins in stoichiometric amounts. (c) 30S subunits are degraded faster than 50S. (d) In total proteome, the quantity of 21 r-proteins decreases during 14 days (short-lived r-proteins) concomitantly with the reduction of cellular RNA. (e) 30 r-proteins are stable and form a pool of free r-proteins (stable r-proteins). Thus r-proteins are present in non-stochiometric amounts in the proteome of *E. coli* during the extended stationary phase.

**IMPORTANCE:** Ribosome degradation has been extensively described from the viewpoint of its main component – rRNA. Here we aim to complement our knowledge by quantitatively analyzing r-protein degradation and stability in the ribosomes as well as in the whole-cell proteome during stationary phase in *E. coli*. R-proteins are considered to be very stable in the proteome. Here we show that a specific set of r-proteins are rapidly degraded after release from the rRNA. The degradation of r-proteins is an intriguing new aspect of r-protein metabolism in bacteria.

## INTRODUCTION

Translation of genetic information into proteins occurs in ribosomes. Ribosome function is vital for the life. Therefore, the quality and quantity of ribosomes in bacterial cells must meet the requirements for protein production during adaptation to changing environmental conditions. Ribosome biogenesis is most active during the early log phase when protein synthesis is at its maximum and is gradually decreasing (1). A single *E. coli* cell contains up to 70 000 ribosomes during the exponential growth on glucose and up to 10 times less after the cessation of growth (1). Ribosome content in bacterial cells is tightly controlled by several regulatory mechanisms during the progression of the growth phase (2-4).

Bacterial ribosomes are composed of three RNA (rRNA) chains and over 50 protein (r-protein) molecules. Proteins make up about one-third of the ribosome by mass, the remaining two-thirds is rRNA. rRNA and r-proteins are organized into two unequal subunits. In *E. coli* the small ribosomal subunit (30S) consists of 16S rRNA (1543 nucleotides) and 21 r-proteins (bS1…bS21). The large ribosomal subunit (50S) comprises of 23S rRNA (2904 nt), 5S rRNA (120 nt), and 33 different protein chains (bL1…bL36). All components are present in a single copy in the mature ribosome (70S) except for the protein bL7/bL12, which is associated with the ribosomes as a tetramer via the uL10 protein (5, 6). The majority of the r-proteins are small and basic. They are important for ribosome assembly and for maintaining ribosome structure and function (6).

Translation initiation starts with the 30S binding the initiator tRNA and mRNA in its decoding center (DC) (7). DC is responsible for the recognition of the appropriate aminoacyl-tRNA (aa-tRNA) anticodon guided by the mRNA codons during the elongation phase of translation (7). The 50S encloses the peptidyltransferase center (PTC), which catalyzes the formation of peptide bonds in the nascent protein chain during elongation as well as the release of the full-length protein at the end of translation (7, 8). Ribosome functional centers DC and PTC are formed of both rRNA and r-proteins (7, 8). Chemical or enzymatic damage of ribosomal components can compromise ribosome functioning (9). Partially active ribosomes must either be degraded or repaired to restore their full functionality (9, 10).

The number of ribosomes per cell starts to decrease during the late log phase due to two processes (11). First, the dilution effect – ribosome synthesis slows significantly while cell division continues. Second, ribosomes are degraded in cells as demonstrated by monitoring rRNA degradation (4). rRNA degradation has been shown to occur under stress conditions e.g. under inhibition of translation in the presence of antibiotics (12), upon rRNA over-production (13, 14), and upon entry into stationary phase (15).

While rRNA degradation is well documented (4, 16), the fate of r-proteins in the stationary phase has remained unknown (17, 18). Here we aim to fill this gap by analyzing the r-protein quantities in the ribosomes as well as the whole-cell proteome during the stationary phase of *E. coli*. We used quantitative mass spectrometry to measure the stability and composition of r-proteins during stationary growth phase of *E. coli* cultures. We show that ribosomes contain nearly all r-proteins in equimolar amounts until late stationary phase, however, the stability of individual r-proteins in proteome is different. Surprisingly, it appears that about two-thirds of 51 r-proteins are stable during stationary growth phase in the course of 14 days while the remaining r-proteins are degraded concomitantly with the rRNA. Thus, r-proteins in *E.coli* proteome are non-stoichiometric during extended stationary phase.

## RESULTS

### Experimental setup

The aim of this study was to determine the fate of ribosomes and r-proteins. In this respect, SILAC (**S**table Isotope **L**abeled **A**mino acids in cell **C**ulture) based experimental approach was used (Fig. 1). *E.coli* cells were grown in MOPS medium supplemented with “heavy” labeled arginine (Arg10) and lysine (Lys8). At the mid-log phase, the culture was further supplemented with a 20-fold molar excess of “light” unlabeled arginine (Arg0) and lysine (Lys0), divided into 8 aliquots, and grown for 14 days. Cell samples were collected at day one (24h), day two (48h), and subsequently in 48h intervals over the following 12 days. Bacterial cell viability analysis has revealed that number of viable cells (CFU) was highest on day two and was subsequently decreasing in modest extent (Figure S1). At each sampling point, four different datasets were generated. **First**, the ribosome particle distribution was analyzed by sucrose gradient centrifugation. Peak areas of the 70S and free ribosomal subunit fractions were quantified and 50S/70S or 30S/70S ratios were calculated. **Second**, the rRNA amount in the samples was estimated, assuming that 80% of the total RNA of bacterial cell corresponds to the rRNA (1, 19). Total RNA was isolated from cells using hot phenol extraction, concentration was measured, and values were normalized against day one. **Third and fourth** respectively, the quantities of r-proteins in the 70S ribosome fraction and the total proteome were determined using SILAC based LC-MS/MS and normalized to the corresponding values of day one. The latter allowed us to directly distinguish r-protein degradation in ribosomes versus in the whole proteome which included the amount of non-bound (‘free’) r-proteins.

**Figure 1.**
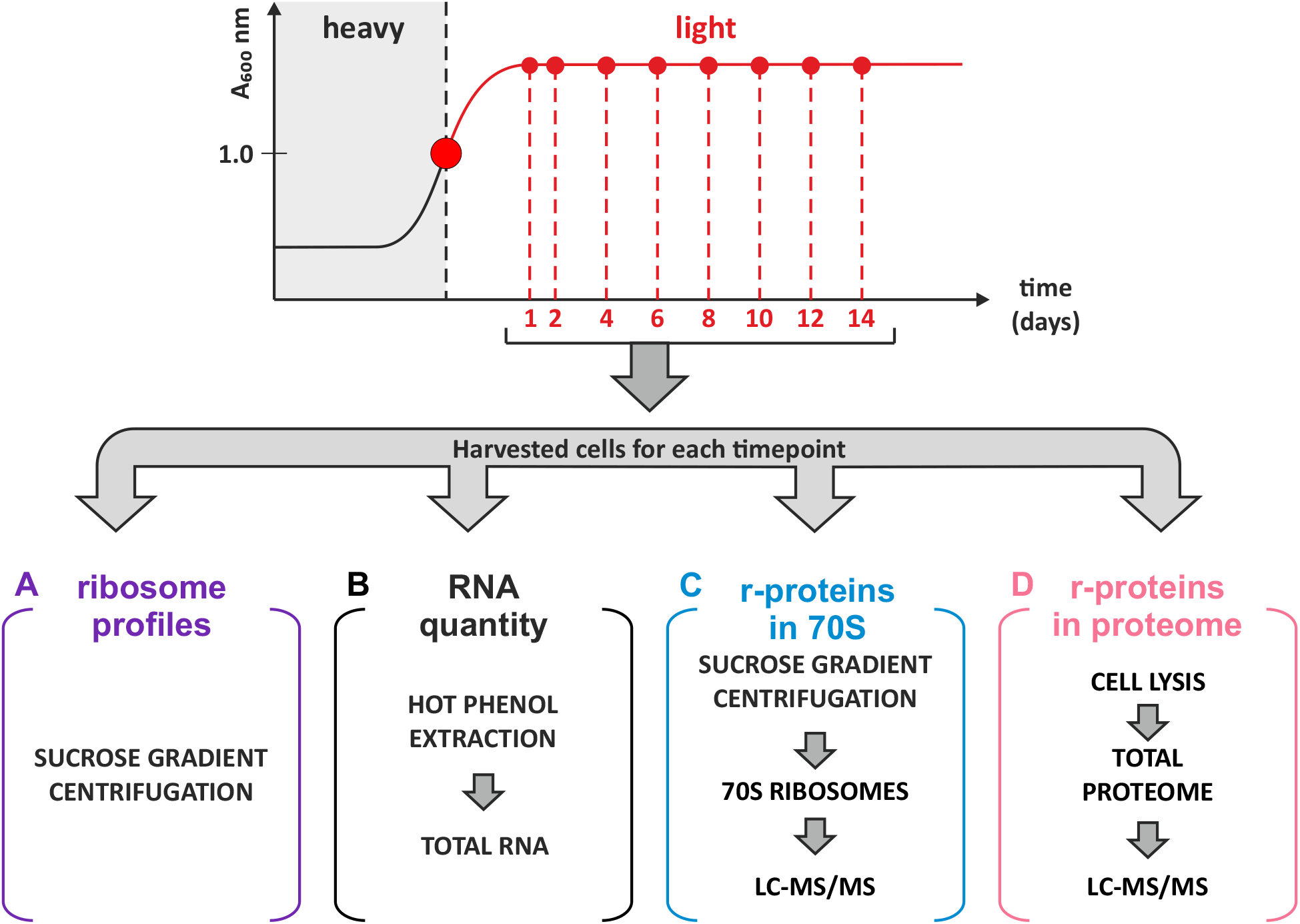
Experimental design. *E. coli* MG-SILAC strain was grown in medium containing heavy labeled arginine and lysine to mid-log phase, followed by a chase with a 20-fold excess of unlabeled (light) arginine and lysine and grown for 2 weeks to obtain stationary phase cells. Samples were collected at time points and divided into three aliquots. (A) Ribosomes were isolated using sucrose gradient centrifugation (purple); (B) RNA quantity was determined by hot-phenol extraction (black); (C) r-protein quantity was determined using LC-MS/MS (cyan); (D) r-protein stoichiometry in total proteome was ascertained using LC-MS/MS (pink).

### Unequal accumulation of ribosome subunits in the stationary phase

To find out how ribosome population changes during the stationary phase, the ribosome content in batch culture over the course of 14 days was analyzed by sucrose gradient centrifugation (Fig.1A). Representative ribosome profiles from day 1, day 10, and day 14 are shown in Fig. 2A (for a complete set of ribosome profiles see Fig. S2). 70S peak from days one to four exhibits a notable shoulder (Fig. 2A and Fig. S2), that was identified as 100S particles according to sucrose gradient analysis using specific conditions as specified in legend of Supplementary Figure S3 (Fig. S3). To quantify how free ribosomal subunit amounts, change comparatively to each other, ratios between peak areas of free subunits and 70S ribosomes (50S/70S and 30S/70S ratios respectively) were calculated from gradient profiles (Fig. 2B). Finally, results were controlled for multiple comparisons using the Bonferroni method (Table S1).

**Figure 2.**
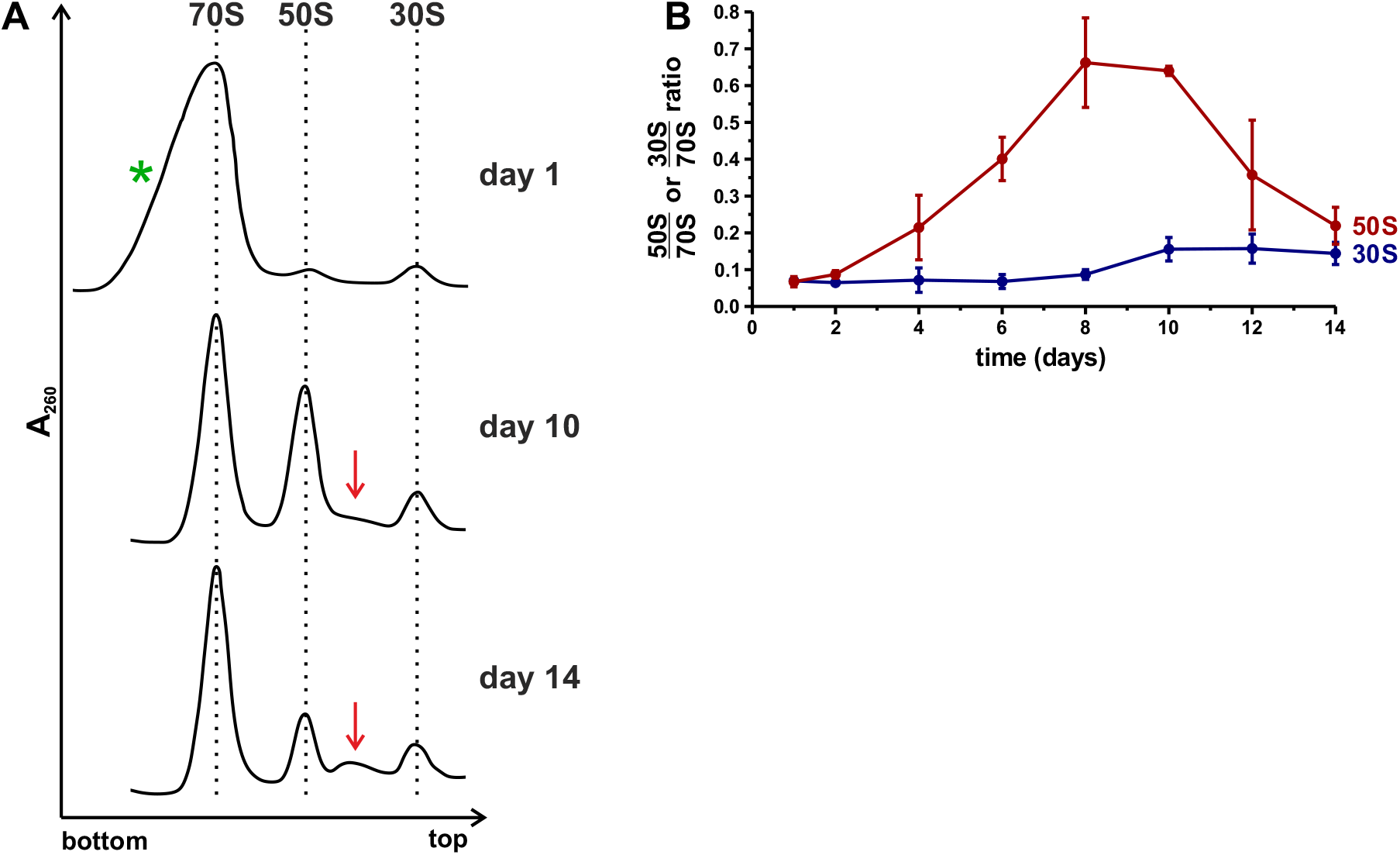
Ribosome profile analysis of stationary phase cells using sucrose gradient centrifugation. Cells were harvested from the stationary phase over the course of 14 days. (A) Representative ribosome profiles from day 1, day 10, and day 14. The direction of sedimentation is from right to left. Red arrows show the accumulation of intermediate particles. The green asterisk indicates notable shoulder present on 70S peak at the beginning of the stationary phase. The complete dataset is shown in Figure S2. The ribosome patterns are normalized to 70S peak. (B) Free subunit to 70S ribosomes ratios calculated based on corresponding ribosome profiles. Areas under the 70S, 50S, and 30S peaks were quantified using ImageJ, subunits/70S ratios were calculated and plotted (Y-axis). Values shown in the figure are the mean of three independent biological experiments with standard deviation (n=3; mean ± SD).

On day one, the 50S/70S and 30S/70S ratios are ≈ 0.07 indicative of low levels of free subunits present in the cells in agreement with published data (20). By day four, the 50S/70S ratio is ≈ 0.21 and further increases up to ≈ 0.64 by day 10, indicating the accumulation of free 50S subunits. In contrast, the 30S/70S ratio remains low over the course of 14 days. After day 10, the 50S/70S ratio decreases down to ≈ 0.19, whereas the 30S/70S ratio remains stable. It is worth noting that after 10 days, particles sedimenting between the 50S and 30S start to accumulate (Fig. 2A and Fig. S2). These 40S-like particles are probably degradation intermediates of the 50S subunits. Accumulation and delayed reduction of free 50S subunits indicates a slower degradation of 50S subunit compared to 30S in the stationary phase.

### Total RNA content decreases in stationary phase

Ribosomes are known to be stable in growing bacteria (21), however, as cells enter the stationary phase, the ribosome amount per cell decreases (15). Based on previous reports that rRNA makes up 80% of the total RNA in cells (1, 19), the rRNA content can be evaluated by quantifying total RNA in cells. To determine the relative quantity of rRNA during stationary phase, the concentration of total RNA in stationary phase cells was measured over the course of 14 days. Total RNA was extracted from 2 mg of wet cell mass, RNA concentration was determined and compared with the measurement from day one. Finally, data were fitted onto a one-phase decay model (Fig. 3) and controlled for multiple comparisons using the Bonferroni method (Table S2). As a control, the amount of total RNA was determined before cells enter stationary phase (3, 6, and 12 hours after the start of the growth). Total RNA concentration per cell mass starts to decrease at the late log phase (Fig. S4), continues with the progression of the stationary phase, and reaches an apparent plateau on day 6 (Fig. 3). After day 6, no statistically significant decrease in RNA concentration was detected. At day 6 the amount of RNA was decreased by 65% compared to day one (Fig. 3). 16S and 23S rRNA were quantified in total RNA by dot-plot hybridization (Fig. S5). The fraction of both 16S and 23S rRNA decreases over time, however, the decrease of 23S rRNA fraction is significantly slower than 16S rRNA on days 6 – 10. This is reminiscent to the free ribosome subunit content (compare Fig. 2B and Fig S5C). In conclusion, the decrease of total RNA content in the cells attributes to the degradation of rRNA in the cells.

**Figure 3.**
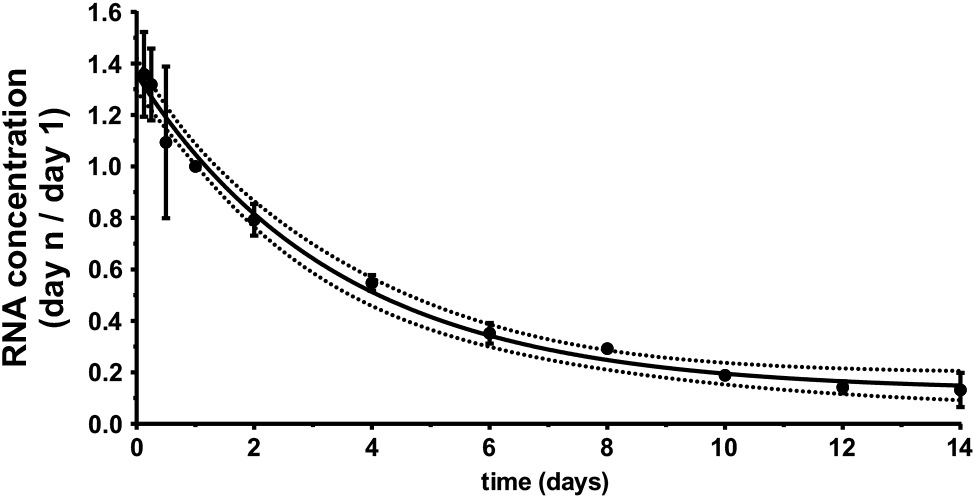
Total RNA content in cells decreases during the stationary phase. Samples were collected over the course of 14 days. Total RNA was extracted from cells using hot-phenol extraction and ethanol precipitation. RNA concentration was determined by measuring absorbance at 260 nm and normalized to the corresponding value on day one (Y-axis). Data were fitted into a one-phase decay model, where black solid line represents a mean and dotted line 95% confidence interval. Values shown in the figure are the mean of three independent biological experiments with standard deviation (n=3; mean ± SD).

### Stationary phase ribosomes contain equimolar amounts of r-proteins

In our previous study, we showed that the ribosome r-protein composition does not change during the transition of the cell culture from the exponential to the stationary phase except for bL31 and bL36 paralogues (22). Also, it has been shown, that ribosomes contain stoichiometric amounts of r-proteins during exponential growth (5, 6, 23). In the present study, we focused on the changes in r-protein stoichiometry in the ribosomes during the stationary phase. 70S ribosomes were isolated from stationary phase cells and mixed in a 1:1 molar ratio with the internal reference – a medium-heavy labeled (Arg6, Lys4) 70S ribosomes isolated from the mid-log growth phase. The addition of internal reference enables the comparison of individual r-protein stoichiometries in ribosomes at different timepoints. Proteins were digested into peptides and analyzed using LC-MS/MS. Based on MS results, the relative quantities of 51 r-proteins in ribosomes were ascertained, while proteins bL31A, bL31B, bL35, bL36A, and bL36B were not quantified due to an insufficient number of unique peptides. Finally, results were controlled for multiple comparisons using the Bonferroni method and only statistically significant results are discussed.

R-protein quantities in the 70S ribosomes from the stationary phase bacteria are presented as (L+H)/M ratio (Fig. 4, complete dataset shown on Fig. S6, and statistical analysis in Table S3). (L+H)/M ratio compares relative quantities of r-proteins in the stationary phase samples (L+H) to the reference ribosomes (M). 50S r-protein (L+H)/M ratios are normalized to the average of (L+H)/M ratio of all 50S r-proteins. Similarly, 30S r-protein (L+H)/M ratios are normalized to the average of (L+H)/M ratio of all 30S r-proteins. For 43 of the analyzed r-proteins, the (L+H)/M ratio of 1 ± 10%, meaning that these r-proteins are present in stoichiometric amounts (Fig. 4, Fig. S6, and Table S3). However, for proteins bS1 and bS21, the (L+H)/M ratio decreases over time. bS1 has been previously found in non-stoichiometric amounts on 70S ribosomes (24-26). bS20 exhibits an (L+H)/M ratio of 0.75 throughout the entire stationary phase. This systematic difference is likely caused by the unbalanced bS20 composition of the medium-heavy labeled reference ribosomes. In the case of bL7/bL12 and uL10, the (L+H)/M ratio is highly variable (Fig. 4, see also Fig. S6, and Table S3). It has been shown that bL7/bL12 and uL10 are loosely bound to the ribosome (27), which explains the variability arising from the partial loss of the aforementioned proteins during ribosome isolation. In case of proteins bL17, uL22, and bS6, the (L+H)/M ratio increases during the stationary phase. However, when isolated 70S ribosomes are further disassociated into 50S and 30S subunits, the stoichiometry of these proteins is restored (Fig. S7). This increase can be attributed to a non-specific binding of bL17, uL22, and bS6 to the 70S ribosomes during isolation (Fig. S7). Loss of proteins bS1 and bS21 demonstrates another level of ribosome heterogeneity during stationary phase in addition to the replacement of paralogous proteins bL31A and bL36A by their respective B paralogs (22). Taken together, for a majority of the r-proteins, their stoichiometry remains constantly unchanged in the 70S ribosomes during the stationary phase.

**Figure 4.**
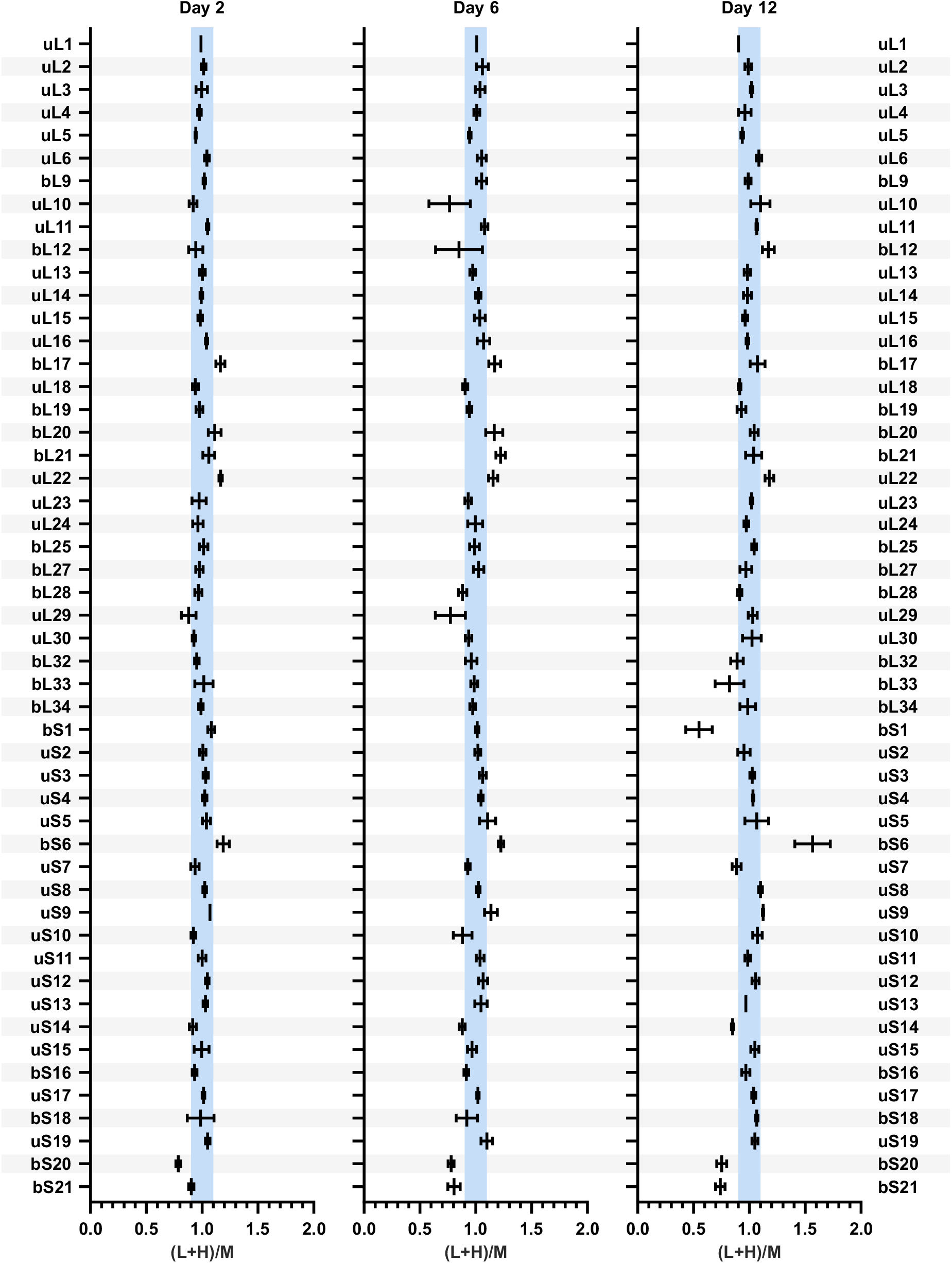
R-protein quantity in the 70S ribosomes during the stationary phase. Cells were collected over the course of 14 days. 70S ribosomes were isolated and mixed in 1:1 ratio with medium-heavy labeled reference 70S ribosomes for r-protein quantification using LC-MS/MS. Proteins bL31A, bL31B, bL35, bL36A, and bL36B were now quantified due to insufficient number of unique peptides. Datasets from days 2, 6, and 12 are shown, the complete set of all datasets are shown in Supplementary figure S6. The relative quantities of 51 r-proteins are presented as the (L+H)/M ratio (L+H = sample; M = reference). 50S r-protein (L+H)/M ratios are normalized against the average of (L+H)/M ratio of all 50S r-proteins. 30S r-protein (L+H)/M ratios are normalized against the average of (L+H)/M ratio of all 30S r-proteins. The blue box marks ±10% range of (L+H)/M ratio. Values shown in the figure are the mean of three independent biological experiments with standard deviation (n=3; mean ± SD).

### R-proteins are present in stoichiometric amounts in early stationary phase proteome

To shed a light into the stoichiometry and stability of r-proteins during prolonged stationary phase we first performed a quantitative analysis of the early stationary phase proteome (day one). The stoichiometry of individual r-proteins in the total proteome on day one was compared to that of r-proteins in the reference 70S ribosomes (Fig. 5A and Table S4). Total protein from day one cells was mixed in 24:1 mass ratio with the medium-heavy labeled 70S ribosomes and analyzed as described above. Forty-eight out of the 51 detected r-proteins exhibits the (L+H)/M ratio of approximately 1 (± 10%). This indicates that they are present in the total proteome of early stationary phase cells in the same stoichiometry as in the ribosomes. R-proteins bL7/bL12 and bS1 had (L+H)/M ratios of 1.3 and 2.0, respectively (Fig. 5A; statistical analysis in Table S4). This means that there are 30% more bL7/bL12 and two-times more bS1 in the cells compared to the ribosomes. This is in a good agreement with earlier data (28, 29), demonstrating that these proteins have significant free pools in vivo. Given that according to our data, most of the r-proteins are present in the same stoichiometry in the early stationary phase total proteome as in the ribosomes, day one proteome dataset was further used as a reference.

**Figure 5.**
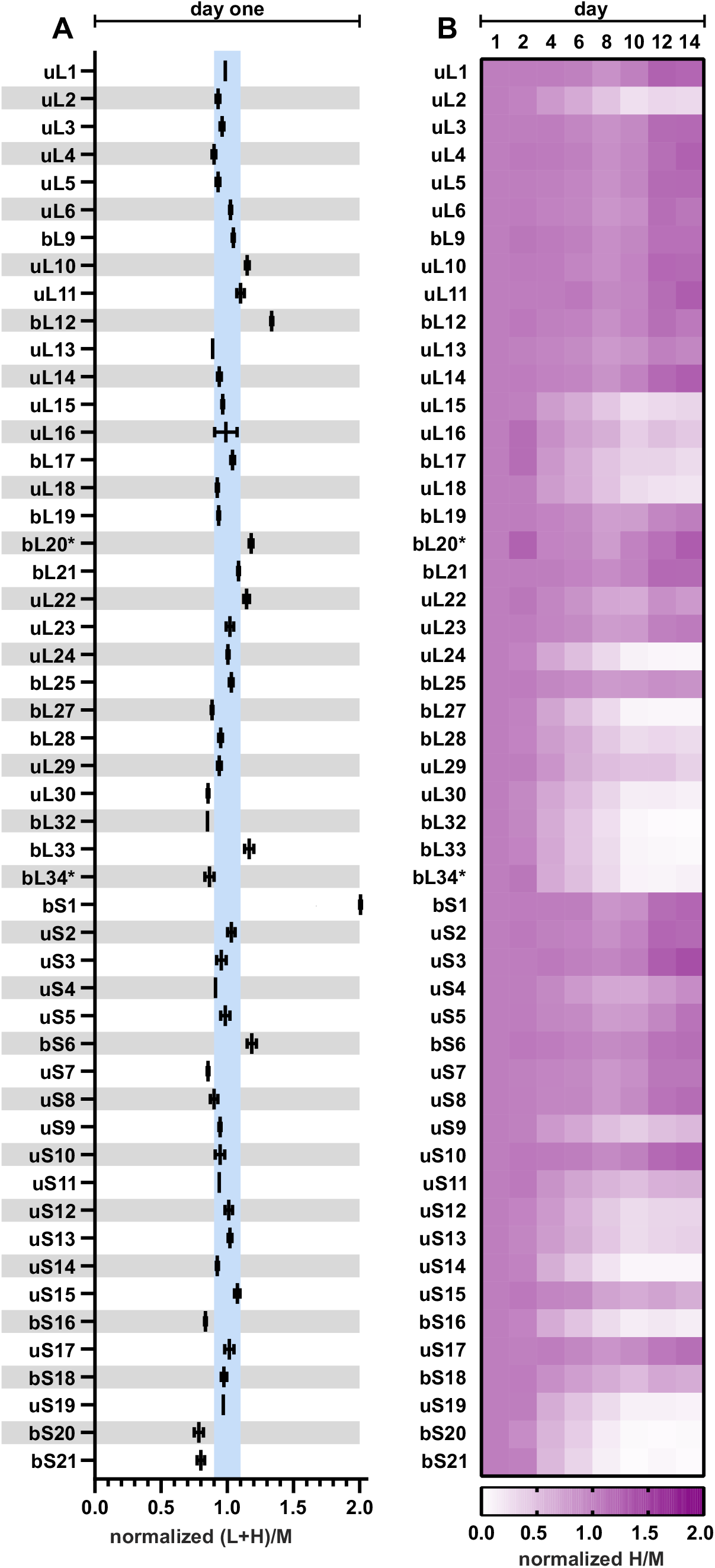
R-proteins become progressively non-stochiometric during stationary phase. Cells were collected over the course of 14 days, lysed, and total protein was mixed in a 1:1 ratio with medium-heavy labeled total protein from the reference cells for r-protein quantification using LC-MS/MS. Proteins bL31A, bL31B, bL35, bL36A, and bL36B were now quantified due to insufficient number of unique peptides. (A) Day one r-protein stoichiometry. R-protein relative quantity is presented as an (L+H)/M ratio (L+H – sample; M – reference). 50S r-protein (L+H)/M ratios are normalized against the average of (L+H)/M ratio of all 50S r-proteins. 30S r-protein (L+H)/M ratios are normalized against the average of (L+H)/M ratio of all 30S r-proteins. The blue box marks ±10% range of (L+H)/M ratio. Values shown in the figure are the mean of two independent biological experiments with standard deviation (n=2; mean ± SD). (B) Heatmap representing r-protein stoichiometry in the total proteome over the 14 days of stationary phase. Normalized H/M ratio describes relative quantities of r-proteins compared to day one. Proteins marked with an asterisk (*) had only one quantified peptide in datasets. Values shown in the figure are the medians of three independent biological experiments (n=3; median).

### R-protein stoichiometry in the total proteome changes during the stationary phase

As we demonstrated that r-protein are stochiometric in early stationary phase, we next wanted to determine if this stoichiometry changes during stationary phase. For that, we analyzed the r-protein quantities in the total proteome during the course of 14 days. The H/M and L/M ratios describe r-protein subpopulations that are synthesized in the early-log phase (H/M) or the late-log and after the entry to stationary phase (L/M). Changes in the 51 r-protein H/M ratios are summarized on the heatmap on Fig. 5B. For 30 r-proteins (uL1, uL3, uL4, uL5, uL6, bL7/bL12, bL9, uL10, uL11, uL13, uL14, bL19, bL20, bL21, uL22, uL23, bL25, bS1, uS2, uS3, uS4, uS5, bS6, uS7, uS8, uS10, uS11, uS15, uS17, and bS18) the H/M (Fig. 5B) and L/M (Fig. S10) ratios do not change in the total proteome. Hence, their relative quantity per total protein mass remained the same as in day one (Fig. 5B; Fig. S8; Fig. S9; Fig. S10; Table S5). Since these r-proteins show no change in measured quantities, they are considered as stable r-proteins. On the other hand, for 21 r-proteins (uL2, uL15, uL16, bL17, uL18, uL24, bL27, bL28, uL29, uL30, bL32, bL33, bL34, uS9, uS12, uS13, uS14, bS16, uS19, bS20, and bS21) the H/M (Fig. 5B) and L/M (Fig. S10) ratio decreases over the next 14 days, meaning they are degraded (Fig. S8; Fig. S9; Table S5). As their measured quantity decreases in stationary phase, they are named short-lived. It is important to note, that both bS1 and bS21 disassociate from ribosomes, but while bS1 remains in the total proteome (stable), the bS21 is rapidly degraded (short-lived). In contrast to the r-protein stoichiometry in the ribosome (Fig. 4 and Fig. S6), the r-proteins are found in non-stoichiometric amounts in the total cell proteome. In conclusion, according to stationary phase proteome data, the r-proteins can be divided into two distinct groups containing: (a) 30 stable r-proteins and (b) 21 short-lived r-proteins.

### R-protein degradation rates are disparate from each other

To quantitatively evaluate the r-protein degradation via their degradation rates, the r-protein dynamics in the stationary phase were modeled using non-linear regression analysis. R-protein quantities in the course of 14 days were fitted into the following models: (a) exponential model – plateau followed by one phase decay; and (b) linear model – a straight line. The fit between data and two models was evaluated using Akaike’s information criterion (30). Two representative examples, one for stable (uL6) and one for short-lived protein (bS16), are presented in Fig. 6 (for the complete dataset see Fig. 3; Fig. S8; Fig. S9). The quantity of uL6 does not change during 14 days following the linear model (stable protein) while the quantity of bS16 fits the exponential model (short-lived protein). For each r-protein, a degradation rate constant was calculated (K-value for the exponential model and slope for the linear model) (Fig. 7; Table S6). These values characterize how fast the relative amount of the specific protein changes in time. The degradation rates of the r-proteins are compared with their mass (Fig. 7). R-proteins with molecular mass over 15kDa are stable, except for uL2 while half of smaller r-proteins are short-lived. Moreover, the degradation trendlines of short-lived proteins tend to coincide with the RNA degradation curve during stationary phase (Fig. 6). At the same time, stable r-protein quantities remain almost unchanged in the total proteome during day 1 to day 14.

**Figure 6.**
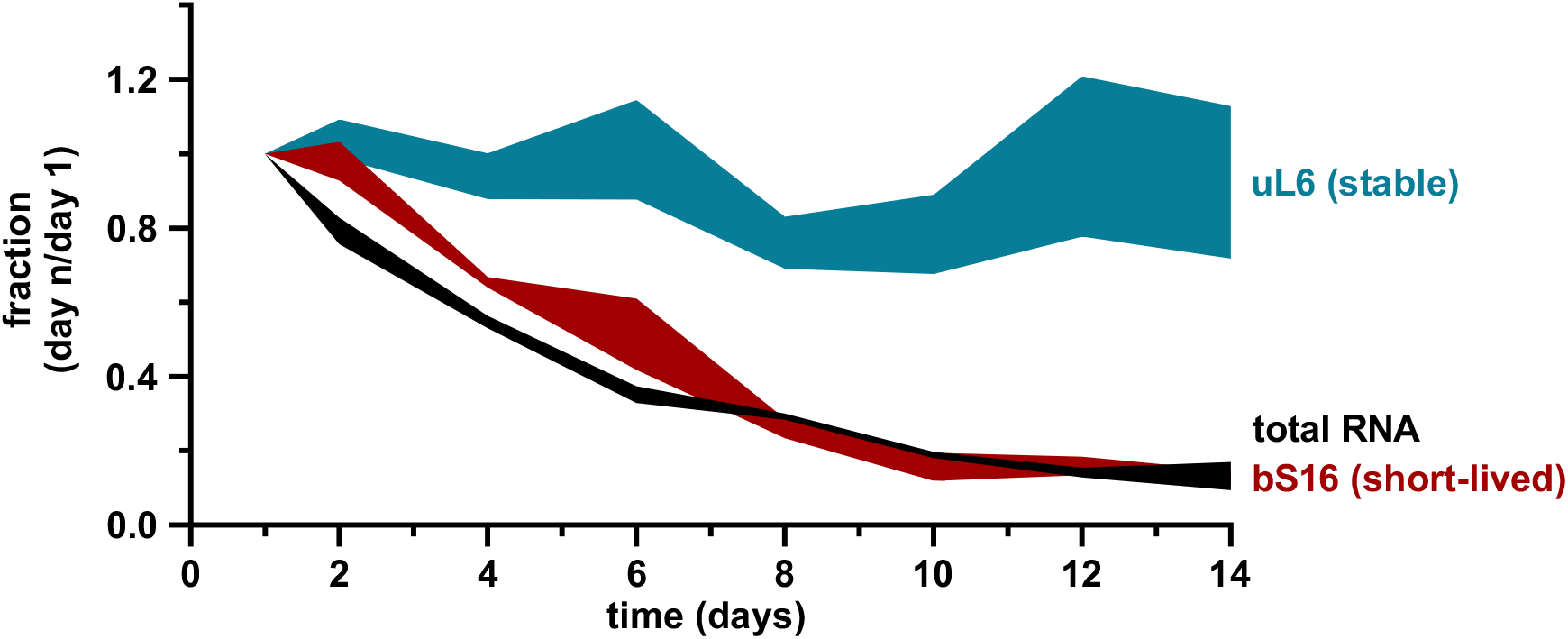
Cellular content of total RNA and r-proteins in stationary phase. Datasets showing quantities of total RNA (black) (Figure 3) and r-protein uL6 (teal) and bS16 (red) (Figure S6 and S7) in stationary phase were compared to each other. Quantity is represented as a fraction from day one during stationary phase (Y-axis). Values shown in the figure are the mean of three independent biological experiments with standard deviation (n=3; mean ± SD).

**Figure 7.**
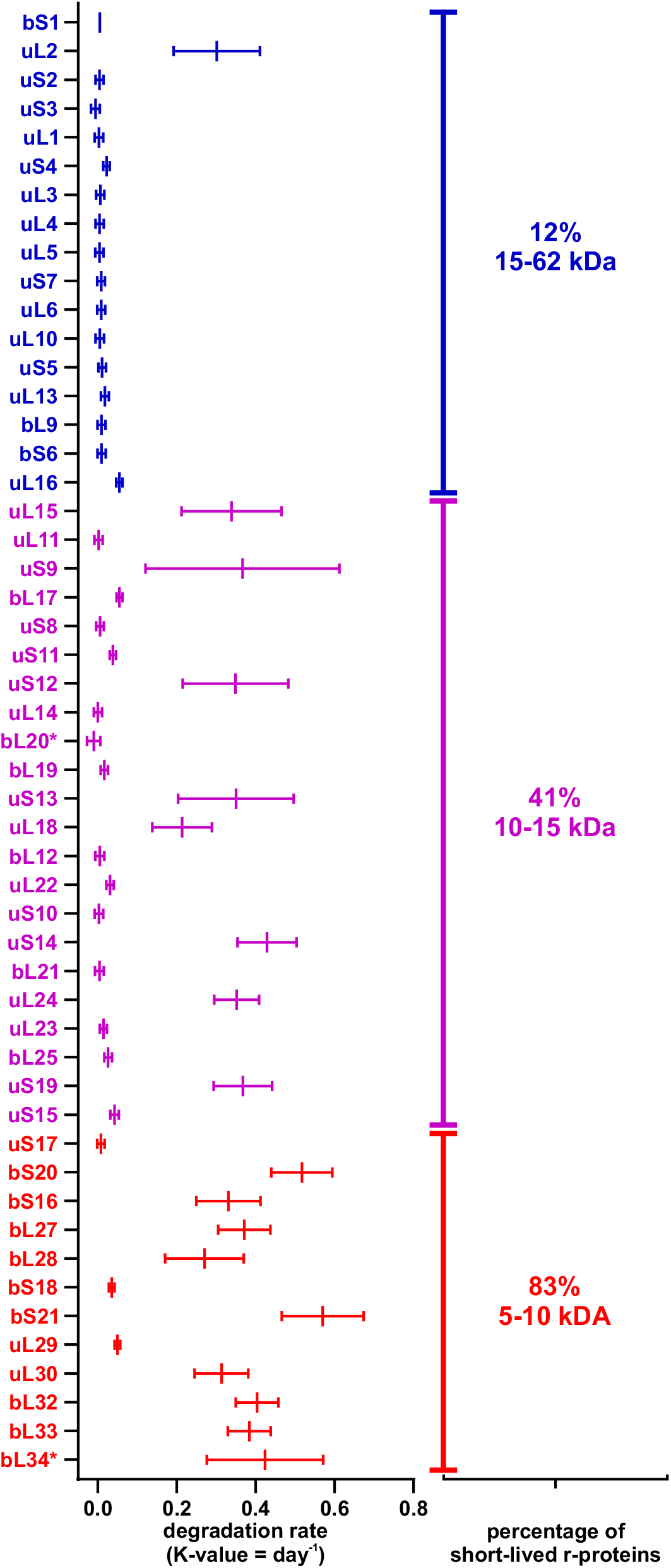
R-protein degradation rate is related to its molecular weight (MW). The r-protein degradation rates were determined using linear regression analysis. X-axis – degradation rates (K-value = day^−1^). Y-axis – r-proteins sorted by molecular weight (from top to bottom). Values shown in the figure are the mean of three independent biological experiments with standard deviation (n=3; mean ± SD).

To sum up, the quantities of stable proteins can be fitted into the straight-line model, while quantities of short-lived proteins can be characterized by the plateau followed by one phase decay model (Fig. 6; Fig. S11; Fig. S12; Table S6). Accordingly, a specific set of r-proteins is not degraded concomitantly with the rRNA, instead forming a free pool of r-proteins in the cell (Table S7).

## DISCUSSION

In this paper, the stability of ribosomes and their components is determined in *E. coli* during the extended stationary phase. Ribosome gradient profiles reveals an accumulation of the free 50S subunits while no significant accumulation of the free 30S subunits is detected during the extended stationary phase (Fig. 2B; Fig. S2). Therefore, free 30S subunits are degraded faster than free 50S subunits, leading to a non-stoichiometric ribosome subunit content during the stationary phase. In accordance, 16S rRNA is degraded faster as compared to 23S rRNA (Fig. S5). However, the decay of 30S subunit rRNA cannot solely account for the loss of the total RNA during the first 6 days of the stationary phase (Fig. 3 and Fig. S5). This is based on the observation that the total RNA content is reduced by 70% but the 30S subunit rRNA constitutes only 30% of the total cellular RNA (1). Therefore, a significant amount of the 50S subunit rRNA (23S and 5S rRNA) must be turned over as well. Indeed, degradation of 16S and 23S rRNA is evident from hybridization experiments (Fig. S5). Again, 23S rRNA appears to be more stable leading to non-stoichiometry of 16S versus 23S rRNA. Ribosome degradation has been suggested to start with the dissociation of ribosome subunit (4), an idea that is supported by the results presented here. An excess of free 30S subunits can lead to non-productive 30S initiation complex formation with initiation factors, initiator tRNA, and mRNA, thereby sequestering mRNA. This in turn can impede translation. Preferential degradation of 30S subunits is likely a failsafe mechanism to avoid the binding and trapping of mRNA molecules in the 30S initiation complexes. We suggest that the degradation of ribosomal subunits likely proceeds via distinct pathways, which is supported by previous studies (12, 31). Delayed degradation of free 50S subunit supports vehement degradation of free 30S subunits (Fig. 2B; Fig. S2). Slower degradation of the 50S subunit is further stressed by the accumulation of the particles sedimenting between the 50S and 30S fractions during the extended stationary phase (indicated by the red arrow on Fig. 2A). In line with ribosome subunit stability is the fact that 23S is more stable as compared to 16S rRNA (Fig. S5). Considering that synthesis of new ribosomes is negligible during the late stationary phase (32), these particles are likely the degradation products of 50S subunits. The fact that no 30S subunits degradation intermediates can be seen on the sucrose gradients further confirms the unbalanced degradation of the ribosome subunits.

Total RNA offers a crude, but effective way to estimate ribosome amount in cells as 80% of it is rRNA (1, 19). Previous studies have shown that ribosomes are degraded as growth conditions of the bacterial culture become limiting (4, 16). In this study, the total RNA levels per cell were measured for 14 days in the stationary phase. Total RNA starts to decrease in the late log and early stationary phase (Fig. S4), which is in a good agreement with previously reported data (15). In this study, we see continuous total RNA decrease after reaching to stationary phase. As stationary phase progresses, the RNA quantity continues to decrease until day 6 when approximately 65% of the total RNA is degraded (Fig. 3). After day 6, no statistically significant decrease in RNA concentration is detected.

Despite extensive ribosome degradation, r-protein quantity in the remaining ribosomes is stable over the course of 14 days in stationary phase (Fig. 4). Two r-proteins, bS1 and bS21, exhibits a significant reduction in the 70S ribosomes (Fig. 4). After 12 days of cultivation, about 50% of the ribosomes have lost bS1, while bS21 is missing from 25% of 70S ribosomes (Fig. 4). Both bS1 and bS21 are important for the initiation step of translation (24, 25, 33). In *E.coli*, bS1 takes part in the recruitment and positioning of mRNA to the 30S to form the initiation complex and it is a part of the translating ribosomes (34). bS1 has been shown to be present at sub-stoichiometric amounts in the non-translating ribosomes (24). Proteins bS1 and bS21 may be removed from the ribosomes as an additional mechanism of regulation of translation initiation during the stationary phase. Moreover, sub-stochiometric r-protein composition is an example of the ribosome heterogeneity in stationary growth phase.

Given that most r-protein stoichiometry in the 70S ribosomes does not change during the stationary phase, we compared this with r-protein stoichiometry in the total cell proteome. During transition from exponential to stationary growth, r-proteins are detected in the total proteome in the same stoichiometry as they are found in ribosomes i.e., one copy of each individual r-protein per ribosome (Fig. 5A). Thus, at this time point, there are no large pools of free r-proteins in the cells except for the proteins bS1 and bL7/bL12 (Fig. 5A). During the progression of the stationary phase, the equimolar content of the r-proteins is maintained in the ribosomes (Fig. 4). Our total proteome analysis clearly demonstrates that the stoichiometry of r-proteins is lost during the stationary phase and a specific set of r-proteins accumulate as a free pool. Thirty out of 51 analyzed r-proteins are found to be stable in course of a 14-day timescale, while 21 r-proteins are a subject of faster degradation (Fig. 5B). Degradation of these r-proteins occurs concomitantly with the diminishing of rRNA, although with a small delay (Fig. 6).

It is evident that r-proteins are stable as long as they are an integral part of the ribosome and become a potential substrate for the degradation machinery after the rRNA is degraded. Considering the differences in accumulation of the free ribosomal subunits, one would expect that the 30S subunit proteins have faster degradation rates. However, according to our experimental data, both subunits contain r-proteins belonging to stable and short-lived r-proteins roughly to the same extent. Most of the smaller-sized r-proteins belong to a short-lived group, while larger-sized r-proteins tend to be stable in stable group. (Fig. 7). A notable exception is the largest r-protein of the 50S subunit, uL2, which is degraded during the stationary phase. The proteins, that are known to form a significant free pool already in the exponential phase (bS1 and bL12) were as stable in our study (28, 29). The proteins with known non-ribosomal function (bS1, uS10, and uL4) (6) are expectedly detected as stable (Fig. 5B). The physiological significance of the other stable r-proteins remains elusive. We cannot exclude the possibility that stable r-proteins have an unknown function(s) during the stationary phase. However, it is highly possible that their stability is not a question of function, but an exception in regards to their structure or conformation in the stationary phase. Previous studies have shown that after rRNA degradation, unbound r-proteins can reintegrate into new ribosomes (35). When cell culture transitions into a new lag phase, these stable r-proteins can be potentially incorporated into newly synthesized ribosomes, allowing faster adaptation of bacterial growth in response to the environmental changes.

In this study, we determined that some r-proteins are not degraded concomitantly with the rRNA, but instead form a free pool of r-proteins in the cell (Table S7). Previously, free pool sizes of r-proteins have been determined in the cell under exponential growth conditions (29). Comparing our results characterizing the stationary growth phase to exponential growth phase, free pool sizes of r-proteins are 20 to 2000 times larger in stationary phase. For example, Chen et al. 2012 reported a free pool size of uL10 was 0.015 ± 0.001 per ribosome. However, in stationary phase the free pool size of uL10 is 1.750 ± 0.533 per ribosome. There are also differences in proteins that form free pools in exponential or stationary phases (Table S7). We conclude, that free pool sizes of r-proteins differ both in quantity and quality between exponential and stationary phase cell cultures.

The fact that r-proteins are present in equimolar quantities makes them a useful internal standard for proteomic analysis of bacteria. However, it should be considered that during the stationary phase, the stoichiometry between individual r-proteins is lost in the cell proteome.

Based on the results reported in this paper, we propose the following model of r-protein dynamics in the stationary phase. RNA content per cell decreases by 65% from day 1 to 6. During the early stages of the stationary phase, free 30S subunits might be degraded faster than free 50S subunits. The latter are degraded predominantly during the later stages of the stationary phase. 21 r-proteins are degraded in the stationary phase alongside rRNA, while 30 r-proteins remain in the cell as a free pool. The stability and potential functionality of these stable r-proteins is an intriguing new aspect of r-proteins and their part in cellular systems.

## METHODS

### Cell growth

*E.coli* MG1655-SILAC (*F-, λ-, rph-1, ΔlysA, ΔargA*) strain was grown in *MOPS* medium (36), supplemented with 0.1 mg/ml “heavy” arginine (Arg10 - [^13^C]_6_H_14_[^15^N]_4_O_2_) and lysine (Lys8 - [^13^C]_6_H_14_[^15^N]_2_O_2_) (SILANTES, Germany). At mid-log (A_600_ ≈ 1), 2 mg/ml of unlabeled arginine (Arg0) and lysine (Lys0) were added to the culture. Cell culture was divided into 8 separate batches and growth was continued for a maximum of 14 days. Cells were harvested by low-speed centrifugation (4500 g/15 min) after 24 and 48 hours (from here on referred to as day one and day two) of growth and subsequently on days 4, 6, 8, 10, 12, and 14. The experiment was carried out in triplicate.

As an internal reference, *E. coli* MG1655-SILAC cells were grown in MOPS medium supplemented with 0.1 mg/ml medium-heavy arginine (Arg6 - [^13^C]_6_H_14_N_4_O_2_) and lysine (Lys4 – C_6_H_10_[^2^H]_4_N_2_O_2_) (SILANTES, Germany). Cells were grown to mid-log (A_600_ ≈ 1) and harvested by low-speed centrifugation (4500 g/15 min).

### Total RNA analysis

Total RNA was extracted from 2 mg of wet cell mass (from the stationary phase) using hot-phenol extraction. Cells were suspended in 200 µl of buffer A (0.5% SDS and 10 mM EDTA). 200 µl of phenol/H_2_O (pH 5.5) was added to the cell suspensions, mixed, and incubated at 65°C for 30 minutes. Samples were centrifuged (16000 g/10 min) and 150 µl of water phase was transferred to a new tube. 150 µl of buffer A was added to the remaining phenol mixture, mixed, and centrifuged (16000 g/10 min). 150 µl of water phase was again moved to a new tube (total of 300 µl). Then, 400 µl of chloroform was added, mixed, and phases were separated by centrifugation (16000 g/10 min). 200 µl of water phase was transferred to a new tube and RNA was sedimented by adding 5 volumes of 96% ethanol and 0.3M of sodium acetate (pH 5.5) and incubating at -20°C overnight. Precipitation was collected by centrifugation (16000 g/10 min), washed 2 times with 96% ethanol, and dried at 37°C for 5 min. RNA precipitate was dissolved in water and absorbance at 260 nm (A_260_) was measured. RNA concentrations (A_260_) were normalized by dividing with the day one A_260_ value (day n / day one). Total RNA concentration values were analyzed across all time points using the two-way analysis of variance (ANOVA) statistical test in software GraphPad 7.0. Statistical values to corresponding experiment are found in Table S2. Total RNA concentration values were also fitted into one-phase decay model in software GraphPad 7.0. One-phase decay model: Y=(Y0 - Plateau)*exp(-K*X) + Plateau, where X is time, Y starts at Y0 and then decays down to plateau with one phase, and K is the rate constant in units that are reciprocal of the X-axis units. For our data plateau and K-values were restricted to being larger than 0.

### Sucrose gradient centrifugation

Cell pellets were suspended in lysis buffer [20 mM Tris (pH 7.5), 100 mM NH_4_Cl, 10 mM Mg-acetate, and 6 mM β-mercaptoethanol]. After the addition of DNase I (40 units/ml), the cells were disrupted with glass beads using Precellys 24 homogenizer (6000 rpm, 4°C, 3×1 min, pause 1 min). Lysate was cleared of cell debris by centrifugation (16000 g, 20 min at 4°C). Maximum of 100 A_260_ units (for precise amounts look Table S8) of supernatant was loaded onto a 15–25% sucrose gradient in OV-10 buffer [20 mM Tris (pH 7.5), 100 mM NH_4_Cl, 0.25 mM EDTA, and 6 mM β-mercaptoethanol] supplemented with 10 mM Mg-acetate and centrifuged at 56000 g for 16 h in a Beckman SW-28 rotor. Ribosome profiles were recorded at 260 nm. Areas under the 70S, 50S, and 30S peaks were quantified by ImageJ and corresponding ratios were calculated. Subunit ratios were analyzed across all time points using the two-way analysis of variance (ANOVA) statistical test in software GraphPad 7.0. Statistical values to corresponding experiment are found in Table S1. Fractions containing 70S were collected for further analysis via liquid-chromatography mass spectrometry (LC-MS/MS).

### Ribosome r-protein content analysis

70S ribosomes from stationary phase and reference cells were mixed in 1:1 molar ratio and precipitated with 10% trichloroacetic acid (TCA) overnight at 4°C. Precipitated proteins were pelleted by centrifugation (16000 g for 60 min) at 4°C, washed twice with 80% ice-cold acetone, and air-dried at 37°C for 5 minutes. All subsequent sample preparations were conducted at room temperature. Proteins were dissolved in 50 µL of 8M urea/2M thiourea solution, reduced for 1 h at 56°C by adding 1 mM dithiothreitol (DTT), and carbamidomethylated with 5 mM chloroacetamide for 1 h in the dark. Proteins were digested with endoproteinase Lys-C (Wako) at an 1:50 enzyme to protein ratio for 4 h. Urea concentration in the solution was reduced by adding 4 vol of 100 mM ammonium bicarbonate (ABC) and peptides were further digested using mass spectrometry grade trypsin (enzyme to protein ratio 1:50) overnight. Enzymes were inactivated by the addition of trifluoroacetic acid (TFA) to 1% final concentration. For LC-MS/MS analysis, peptides were desalted on self-made reverse-phase C18 StageTips columns and analyzed by LC-MS/MS using LTQ-Orbitrap XL (Thermo Scientific) coupled with an Agilent 1200 nanoflow LC via nanoelectrospray ion source (Proxeon). 1 mg of purified peptides were injected at a flow rate of 700 nl/min into 75 mm x 150 mm fused silica emitter (Proxeon), packed in-house with Reprosil-Pur 120C18-AQ, 3 mm stationary phase beads (Dr. Maisch GmbH), and eluted over 120 min using linear gradient of 3% to 40% of solvent B (80% acetonitrile and 0.5% acetic acid) in solvent A (0.5% acetic acid) at a flow rate of 250 nl/min. The LTQ-Orbitrap was operated in a data-dependent mode and a „lock mass” option was enabled for m/z 445.120030 to improve mass accuracy. Precursor ion full scan spectra (m/z 300 to 1800) were acquired in the Orbitrap in profile with a resolution 60000 at m/z 400 (target value of 1000 000 ions and maximum injection time 500 msec). The five most intense ions were fragmented in linear ion trap by collision-induced dissociation (normalized collision energy 35.0%) and spectra were acquired in centroid (target value of 5000 ions and maximum injection time 150 msec). Dynamic exclusion option was enabled (exclusion duration 120 s) and ions with unassigned charge state as well as singly charged ions were rejected. The number of peptides per protein analyzed is shown in Table S9).

### Total proteome analysis

Cells were suspended in 10 volumes of 4% SDS, 100 mM Tris-HCl pH 7.5, and 100 mM DTT containing lysis buffer. Cell suspensions were heated at 95°C for 5 min and lysed by sonication (Bandelin) (60 × 1-sec pulses at 50% intensity). Cell debris was removed by centrifugation at 14 000 g for 10 min. Protein concentration was determined at A_280_ using bovine serum albumin (BSA) as a standard. 7.5 µg of total protein from stationary phase cell lysates was mixed in 1:1 ratio with total protein from reference cell lysates. For r-protein stoichiometry in the early stationary phase total proteome, 18 µg of total protein from stationary phase cell lysates was mixed in 24:1 ratio with 70S ribosomes (0.763 µg). Samples were precipitated with 2:1:3 volume methanol:chloroform:water. Protein pellets were suspended in 25 μl of 7 M urea, 2 M thiourea, followed by disulfide reduction with 5 mM DTT for 30 min and cysteine alkylation with 10 mM chloroacetamide for 30 min at room temperature. Proteins were digested with endoproteinase Lys-C (Wako) at an 1:50 enzyme to protein ratio for 4 h. Urea concentration in the solution was reduced by adding 4 vol of 100 mM ABC and peptides were further digested using mass spectrometry grade trypsin (enzyme to protein ratio 1:50) overnight. Enzymes were inactivated by the addition of trifluoroacetic acid (TFA) to 1% final concentration. Peptides were desalted with self-made reverse-phase C18 StageTips columns. The resulting peptides (Table S10) were fractionated and analyzed by LC-MS/MS (37).

### Mass spectrometry data analysis

Data analysis was performed using Maxquant (v1.5.6.0) with default settings (38), except that the minimal peptide length for the specific and non-specific search was 5 amino acids. Unique peptides were used for quantification, main search peptide tolerance was 8 ppm, and variable modification was used for quantitation of oxidation (methionine). The peptide identification search was carried out against *E. coli* K-12 MG1655 protein sequence database from UniprotKB (as of Oct. 2019). The search results were filtered and transformed using Perseus (v1.6.14.0) (39). For proteins bL20, bL33, bL34, bS20, and bS21 MS data analysis was done using the Mascot search engine and Skyline as described in (22). Each protein was quantified through SILAC ratios H/M, L/M, and/or (L+H)/M, comparing unlabeled (L) and/or “heavy”-labeled (H) relative quantities against medium-heavy labeled (M) internal reference. Proteins bL31A, bL31B, bL35, bL36A, and bL36B could not be quantified reproducibly in our datasets due to a small number of unique peptides. H/M, L/M, and (L+H)/M values were analyzed across all time points using the two-way analysis of variance (ANOVA) statistical test in software GraphPad 7.0. Statistical values to corresponding experiments are found in Tables S4 to S6.

### R-protein stability modeling in total proteome datasets

Individual r-protein degradation dynamics were analyzed using non-linear regression in software GraphPad 7.0. Data were fitted into plateau followed by one phase decay model: Y= IF(X<X0, Y0, Plateau+(Y0-Plateau)*exp(-K*(X-X0))), where X is time, Y is Y0 until X=X0 and then decays down to plateau with one phase, and K is the rate constant in units that are reciprocal of the X-axis units. For our data plateau and K-values were restricted to being larger than 0. The alternative model was a straight line: Y= YIntercept + Slope*X. Data were weighted by 1/Y^2^, this minimizes the sum of the squares of the relative distance of points from the curve. The fit of models was compared using Akaike’s information criteria (30). Statistical values to the corresponding experiment are found in Table S6.

## ACKNOWLEDGEMENTS

We are very grateful to Silva Lilleorg, Tiina Tamm, and Margus Leppik for useful discussions and comments on an early draft of the manuscript. We also thank the Proteomics Core Facility of the University of Tartu for their assistance during data collection.

This research was supported by the University of Tartu ASTRA Project PER ASPERA, financed by the European Regional Development Fund (to KR), and by the Estonian Research Council grant PUT 1179 (to JR).

## Data availability

Proteomics data can be found in the EMBL-EBI PRoteomics IDEntification database (PRIDE). Extended supplementary materials can be found in the DataDOI database http://dx.doi.org/10.23673/re-310.

